# Conditional silencing of H-2D^b^ class I molecule expression on dendritic cells modulates the protective and pathogenic kinetics of virus-antigen specific CD8 T cell responses during Theiler’s Virus infection

**DOI:** 10.1101/632265

**Authors:** Zachariah P. Tritz, Robin C. Orozco, Courtney S. Malo, Lila T Yokanovich, Katayoun Ayasoufi, Cori E. Fain, Roman H. Khadka, Megan L. Settell, Mike J. Hansen, Fang Jin, Aaron J Johnson

## Abstract

Theiler’s murine encephalomyelitis virus (TMEV) infection of the central nervous system is rapidly cleared in C57BL/6 mice by an anti-viral CD8 T cell response restricted by the MHC class I molecule, H-2D^b^. While the CD8 T cell response against neurotropic viruses is well characterized, the identity and function of the antigen presenting cell(s) involved in this process is(are) less well defined. To address this gap in knowledge, we developed a novel C57BL/6 H-2D^b^ conditional knockout mouse that expresses an H-2D^b^ transgene in which the transmembrane domain locus is flanked by LoxP sites. We crossed these H-2D^b^ LoxP mice with MHC class I-deficient mice expressing Cre-recombinase under either the CD11c or LysM promoter in order to silence H-2D^b^ restricted antigen presentation predominantly in dendritic cells or macrophages, respectively. Upon challenge with intracranial TMEV infection, we observe that CD11c+ APCs are critical for early priming of CD8 T cells against the immunodominant TMEV peptide VP2121-130 presented in the context of the H-2D^b^ molecule. This stands in stark contrast to later time points post TMEV infection where CD11c+ APCs appear dispensable for the activation of antigen-specific T cells; the functionality of these late-arising antiviral CD8 T cells is reflected in the restoration of viral control at later time points. These late-arising CD8 T cells also retain their capacity to induce blood-brain barrier disruption. In contrast, when H-2D^b^ restricted antigen presentation was selectively silenced in LysM+ APCs there was no overt impact on the priming of D^b^:VP2121-130 epitope-specific CD8 T cells, although a modest reduction in immune cell entry into the CNS was observed. This work establishes a model system which enables critical dissection of MHC class I restricted antigen presentation to T cells, revealing cell specific and temporal features involved in the generation of antiviral CD8 T cell responses. Employing this novel system, we established CD11c+ cells as a pivotal driver of acute, but not later-arising, antiviral CD8 T cell responses against the TMEV immunodominant epitope VP2121-130, with functional implications both for T cell-mediated viral control and immunopathology.

## INTRODUCTION

Theiler’s murine encephalomyelitis virus (TMEV) is a neurotropic picornavirus which has classically been studied as a murine model of multiple sclerosis [1]. Strains of mice susceptible to Daniel’s strain virus, such as SJL mice, experience chronic infection and demyelination of the spinal cord. Meanwhile, resistant mouse strains, including C57BL/6 mice, present with acute encephalitis and seizures before clearing TMEV infection [2-4]. Resistance to TMEV has been linked to the H-2D MHC class I molecules, with various alleles providing different levels of protection [5-7]. One such MHC class I allele, H-2D^b^, is expressed in C57BL/6 mice and is capable of presenting the immunodominant viral peptide, VP2_121-130_, an essential nine amino acid section of the VP2 capsid protein, against which a particularly robust CD8 T cell response is generated [8-12]. This epitope, and the cytotoxic T lymphocyte (CTL) response raised against it, are responsible for the clearance of the Daniel’s strain of TMEV. The involvement of a singular MHC allele in generating productive T cell responses against TMEV makes this virus an ideal model pathogen for interrogating CD8 T cell priming in the brain [13]. While the dynamics and consequences of the CD8 T cell response in this model infection of the central nervous system (CNS) have been well characterized, the identity of the antigen-presenting cell(s) (APCs) responsible for the initial priming of this response remains undefined.

While the infiltration of antigen-specific CD8 T cells into the brain parenchyma in the context of CNS infection, such as West Nile Virus, is well accepted, the specific antigen-presenting cell type(s) required to prime antiviral CD8 T cells has not been fully defined [14-16]. Classically, immune responses against peripheral pathogens are primed by dendritic cells (DCs) [17]. However, this cellular population is largely, but not entirely, absent from the brain parenchyma in the steady state [18-24]. There is evidence that the minimal population of CNS resident DCs can be further augmented by additional DCs migrating into the brain during periods of inflammation [25, 26]. In contrast, macrophages (MΦs) in the perivascular spaces and microglia in the parenchyma are, at least numerically, the predominant APCs within the CNS. Microglia are best understood in regards to their homeostatic role in the brain as a resident phagocyte. Additionally, they have been shown to have some activity as APCs and to upregulate MHC class I and costimulatory molecules during periods of inflammation [27, 28]. MΦs are also capable of presenting antigen to T cells *in vivo*, and the observation of myelin-filled MΦs at the site of multiple sclerosis lesions has raised interest in these APCS as having potential involvement in the priming of autoimmune CD8 T cells [29-32]. Clarifying the roles these three APCs play, either singularly or in concert, in orchestrating CD8 T cell responses against antigens within the CNS would shed light on APC dynamics in this immune-specialized organ.

We sought to address the gap in knowledge regarding the capacity for DCs and MΦs to prime CD8 T cell responses against viral pathogens in the CNS using a novel transgenic mouse model generated by our research team. We introduced a floxed H-2D^b^ transgene into H-2D^b^−/− H-2K^b^−/− (MHC class I deficient) C57BL/6 mice, leaving H-2D^b^ as its sole MHC class I molecule. This provides a platform by which one can engineer ablation of class I restricted antigen presentation capabilities in specific cell types. In the present study, we have crossed this D^b^ LoxP transgenic mouse with mice expressing Cre-recombinase under the CD11c promoter or the LysM promoter. This enabled us to define the role of CD11c+ or LysM+ APCs in the priming of CD8 T cells specific against the D^b^ restricted immunodominant TMEV peptide, VP2_121-130_. Our findings demonstrate a critical role for CD11c+ cells in priming acute CD8 T cell responses against CNS pathogens, despite the relative rarity of these cells within that particular organ. Further, this study is the first to describe a delayed CD8 T cell priming event mediated through APCs in the absence of DC MHC class I. This previously uncharacterized modality of T cell priming might be crucial for long term immunity against brain pathogens and warrants further investigation.

## RESULTS

### Development of C57BL/6 H-2D^b^ LoxP Mice

Class I conditional knockout mice were developed through construction of an H-2D^b^ transgene, in which the transmembrane (TM) domain is flanked by LoxP sites. This transgene was introduced into C57BL/6 mice deficient in H-2D^b^ and H-2K^b^, leaving the floxed D^b^ as the only MHC class I molecule expressed by nucleated cells (Fig 1A) [33]. Crossing this transgenic mouse line with mice expressing Cre-recombinase under the CD11c, LysM, or CMV promoter, also on total MHC class I knockout backgrounds, allowed for CD11c+ cell specific, LysM+ cell specific, or ubiquitous ablation of MHC-I expression, respectively (Fig 1B, Fig 1C, Fig 1D). Resultant mice will be referred to as CD11c D^b^ conditional knockout (cKO), LysM D^b^ cKO, CMV D^b^ cKO, or Cre- littermates depending on whether the F1 generation pups inherited the cre-recombinase gene from their parent. Both LysM and CD11c driven cre-recombinase models have been well characterized by other groups [34]. Although they have some off-target specificities, they have been found to primarily target MΦ and cDCs, respectively [34]. One potential concern in any MHC class I-deficient model is that natural killer cell mediated cytotoxicity through missing-self recognition might lead to a depletion of that cell population. However, we did not detect a change in the percentage of CD11c+ cells within the spleens of unchallenged CD11c D^b^ cKO mice, nor of F4/80+ cells within LysM D^b^ cKO mice, relative to Cre- littermates (Fig 1E).

**Fig 1:**
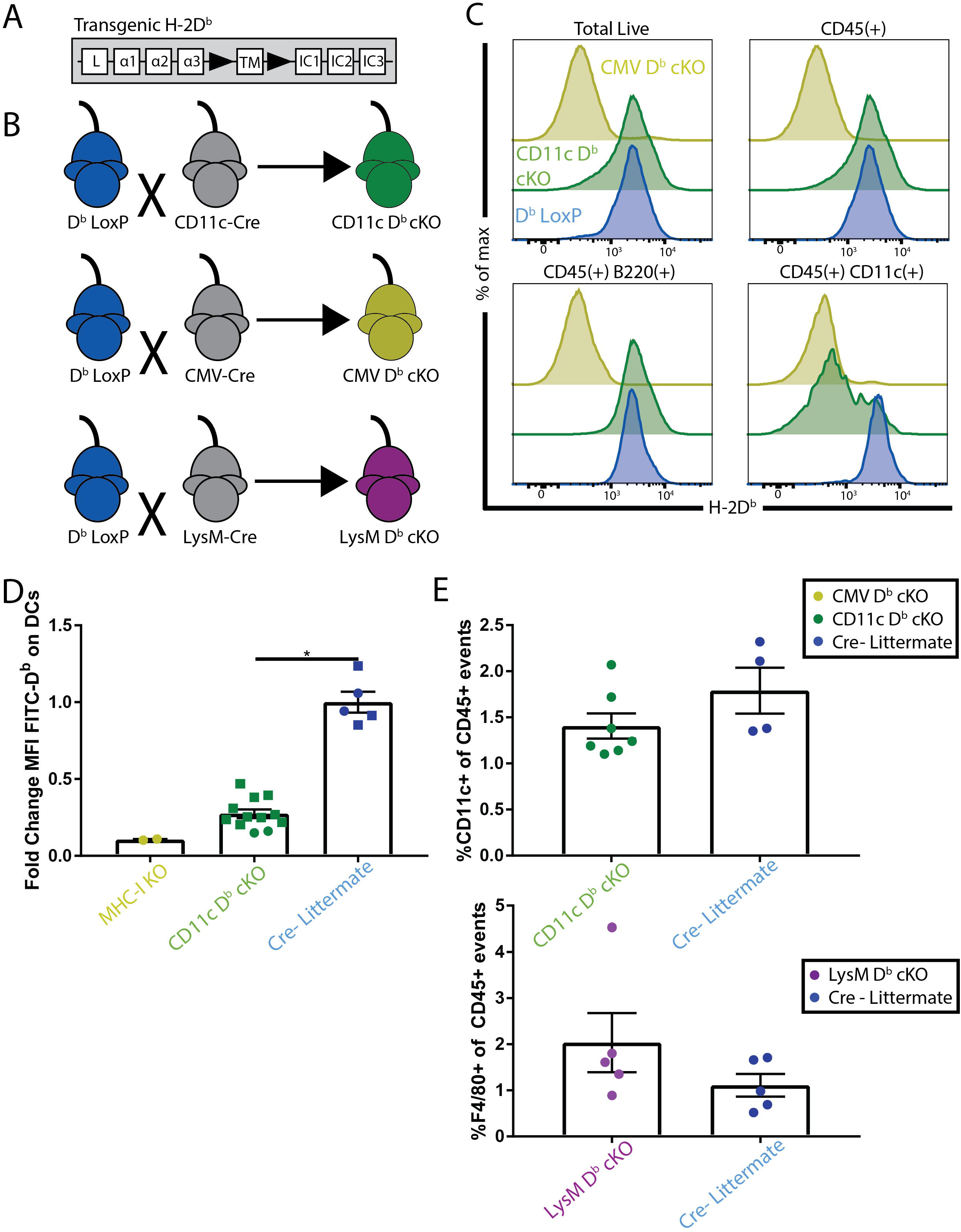
Generation of a mouse model with CD11c+ or LysM+ cell specific H-2D^b^ class I molecule excision. (A) LoxP sites (black arrows) were inserted flanking the transmembrane domain of the H-2D^b^ gene. (B) MHC class I deficient mice expressing Cre-recombinase under the CD11c, LysM or CMV promoter were crossed to D^b^ transgene expressing mice that were otherwise MHC-I deficient. (C) Representative flow plots of live splenocyte subsets from mice seven day post TMEV infection show restriction of CD11c-cre to appropriate cell population. (D) Fold change of median fluorescence intensity (MFI) of FITC-conjugated anti-H-2D^b^ antibody on CD45+CD11c+CD11b^low^F4/80^low^ cells isolated from uninfected spleens. Mann-Whitney Test between CD11c Db cKO (N=12) and Cre- Littermates (N=5). Data taken from two independent experiments with each data point standardized to the average of Cre- Littermates for that experiment. (E) Unchanged percentage (p=0.12) of CD11c+ cells among CD45+ live events in uninfected spleens of CD11c D^b^ cKO animals (N=7) relative to Cre- Littermates (N=4) as assessed by Mann-Whitney test. Unchanged percentage (p=0.31) of F4/80+ cells among CD45+ live events in uninfected spleens of LysM D^b^ cKO animals (N=5) relative to Cre- Littermates (N=5) as assessed by Mann-Whitney test. Data are presented as mean ± SEM. *, p<0.05.

### Conditional deletion of H-2D^b^ in CD11c+ cells results in no steady state immune abnormality

Before investigating the immunologic role of H-2D^b^ on APC subsets during intracranial picornavirus infection, we evaluated the potential of our cre-recombinase system to have overt off-target effects on immune system development. Since DCs of both cDC1 and cDC2 lineages have been reported to contribute to thymic selection and the shaping of peripheral T cell repertoires, we sought to ensure that our DC-specific H-2D^b^ depletion strategy did not disrupt thymocyte development. CD11c D^b^ cKO and Cre- littermates presented equivalent levels of H-2D^b^ on total thymic cells [35]. The only discernible difference in the levels of thymocytes across any developmental stage was that observed in the negative control MHC class I total KO animals, whose lack of CD8 T cells is an expected effect of ubiquitous MHC-I deficiency (Fig 2A, Fig 2B, Fig 2C). Both MΦs and DCs are prevalent in secondary lymphoid organs, so we assessed the TCR Vβ repertoire expressed by the mature splenic T cell pool of naïve animals. As shown in Figure 3, we found no difference in the frequency of the Vβ genes employed by CD8 T cells of either LysM D^b^ cKO or CD11c D^b^ cKO animals relative to Cre- littermates, demonstrating that there were no detectable differences in the TCR repertoire of these transgenic animals (Fig 3A, Fig 3B). These data demonstrate that the novel H-2D^b^ cKO animals have normal T cell development.

**Fig 2:**
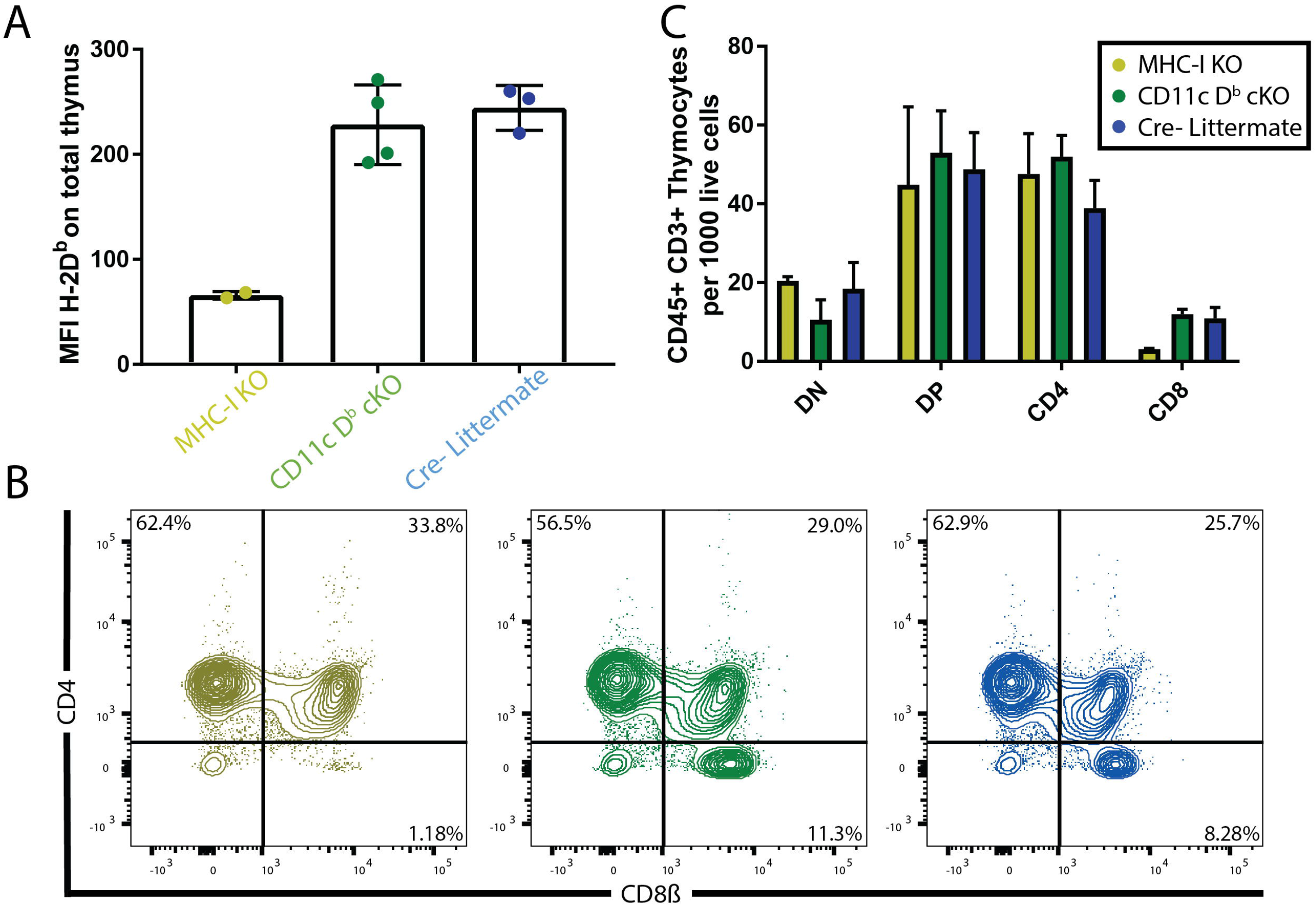
Conditional ablation of H-2D^b^ in DCs does not alter normal CD8 T cell development. (A) Unchanged median fluorescence intensity (MFI) of FITC-conjugated anti-H-2D^b^ antibody on total live cells from uninfected thymi between CD11c D^b^ cKO (N=4) and Cre-Littermates (N=3) (p=0.63) suggests no overt changes in MHC class I expression. Comparison made by Mann-Whitney test. (B) Representative flow plots gated on CD45+CD3+ events within uninfected thymus show normal T cell development in CD11c D^b^ cKO animals. (C) Quantification of live CD45+CD3+ events, either CD4-CD8-(DN), CD4+CD8+ (DP), CD4+CD8-(CD4), or CD4-CD8+ (CD8), within uninfected thymi per 1,000 live events. Comparisons of CD11c D^b^ cKO (N=3) and Cre- Littermates (N=4) were made against MHC-I KO mice (N=2) by 2-way ANOVA with multiple comparisons. Data are presented as mean ± SEM. *, p<0.05.

**Fig 3:**
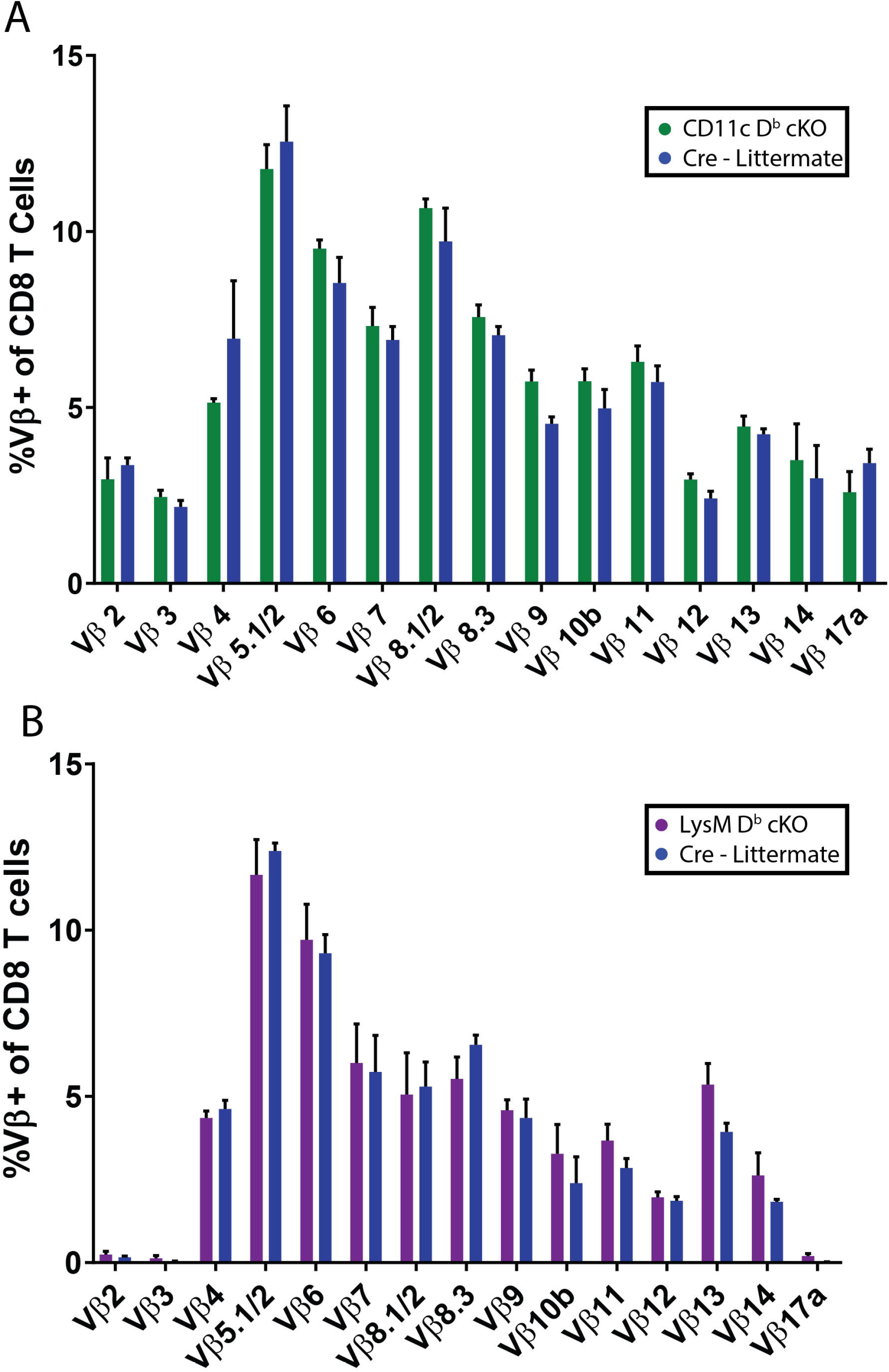
Loss of H-2D^b^ on CD11c+ or LysM+ cells does not impact the Vβ repertoire of naïve splenic T cells. (A) Comparison of the usage of variable regions of TCRβ on CD45+TCRβ+CD8+ gated events from the spleens of uninfected CD11c D^b^ cKO mice (N=4) shows no gross changes relative to the variable region usage observed in Cre- Littermates (N=4). Comparison made by 2-way ANOVA (p=0.52). (B) Comparison of the usage of variable regions of TCRβ on CD45+TCRβ+CD8+ gated events from the spleens of uninfected LysM D^b^ cKO mice (N=5) shows no gross changes relative to the variable region usage observed in Cre- Littermates (N=4). Comparison made by 2-way ANOVA (p=0.96). Data are presented as mean ± SEM. *, p<0.05.

We then assessed H-2D^b^ expression during viral infection, since microglia are known to upregulate CD11c when activated -- an event which would occur during acute TMEV infection. Seven days following intracranial injection with TMEV, flow cytometric analysis of brain tissue revealed no overt changes in expression of H-2D^b^ on the CD11c+ subset of microglia (CD45^intermediate^CD11b+ cells) between CD11c D^b^ cKO and Cre- littermates, suggesting that no unexpected excision of the MHC class I molecule had occurred (Fig 4A, Fig 4B). These findings together establish a robust model for the cell specific deletion of MHC class I on CD11c+ and LysM+ APCs in the periphery with no gross systemic consequences in the steady state.

**Fig 4:**
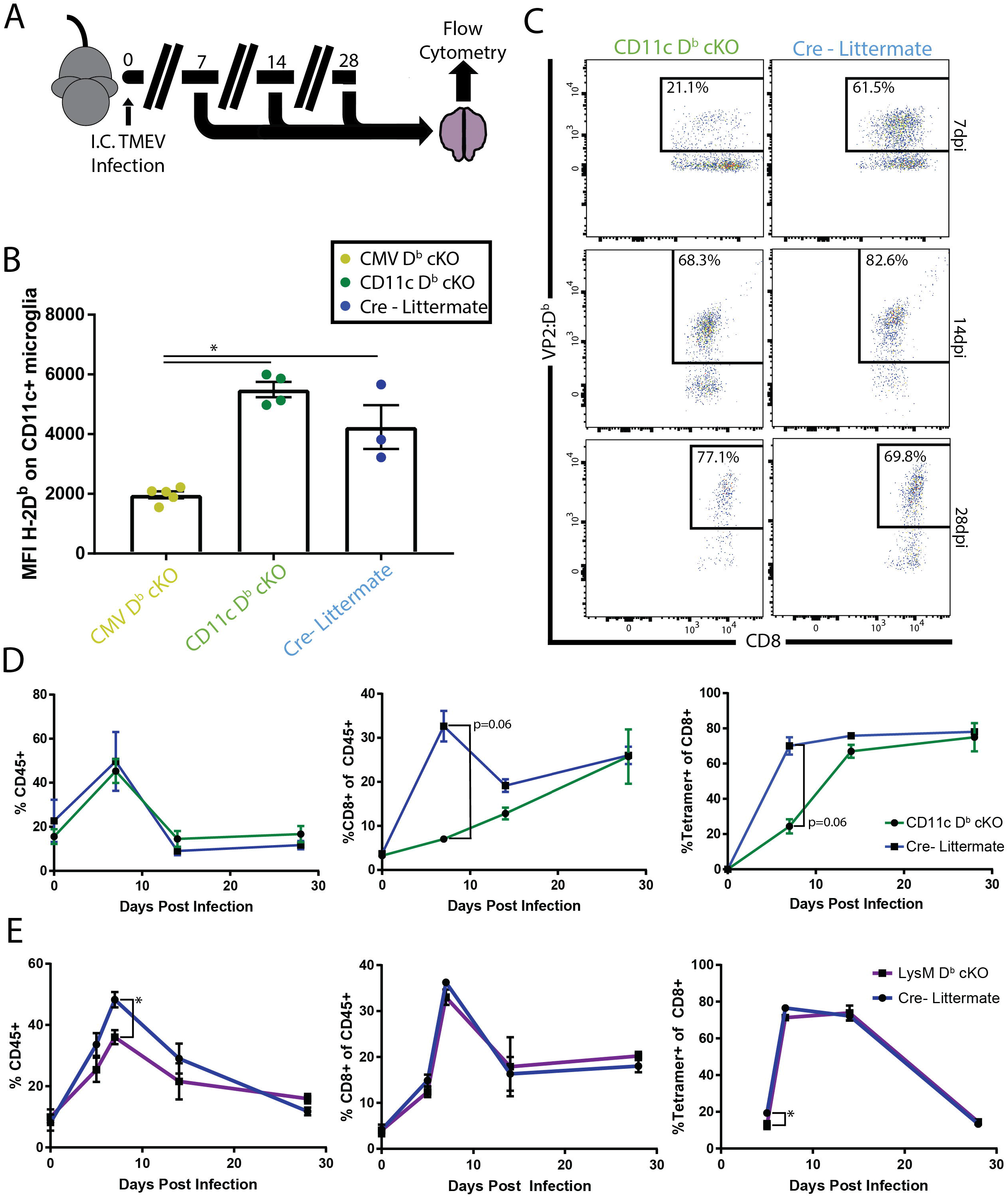
Priming of virus-specific CD8 T cells is impaired in mice with H-2D^b^ deficient CD11c+ cells. (A) Experimental schematic for assessing virus-specific T cells following intracranial TMEV infection. (B) MFI of H-2D^b^ on live CD11c+ microglia (CD45^intermediate^CD11b+) seven days post-infection with TMEV shows no off-target cre activity in CD11c D^b^ cKO animals (N=4) relative to Cre- Littermates (N=3). Levels of H-2D^b^ were compared to CMV D^b^ cKO animals (N=5) as a negative control by Kruskal-Wallis test. (C) Representative flow cytometry data for D^b^:VP2_121-130_ Tetramer-bound live CD45+TCRβ+CD8+ events from brain tissue across the time course of TMEV infection shows profound defect in early T cell priming in CD11c D^b^ cKO animals. (D) Quantitation of flow cytometry data across a time course of intracranial TMEV infection in CD11c D^b^ cKO animals reveals defect in antiviral T cell priming. N>3 for each genotype at each timepoint. Groups were of consistent gender and age within the same timepoint. Comparisons made with Mann-Whitney test between CD11c D^b^ cKO and Cre- Littermates at each time point. (E) Quantitation of flow cytometry data across a time course of intracranial TMEV infection in LysM D^b^ cKO animals reveals overall deficiency in immune cell infiltration of CNS. N>3 for each genotype at each timepoint. Groups were of a consistent gender and age within the same timepoint. Comparisons were made between LysM D^b^ cKO and Cre- Littermates at each time point with Mann-Whitney test. Data are presented as mean ± SEM. *, p<0.05.

### CD11c+ cells are critical for rapid priming of CNS infiltrating antiviral CD8 T cells

We next evaluated the consequence of selective ablation of MHC-I restricted antigen presentation ability in the context of T cell priming against viral infections of the CNS. LysM D^b^ cKO, CD11c D^b^ cKO, and Cre- littermates were infected intracranially with Daniel’s strain TMEV and euthanized at timepoints representative of both acute and chronic infection (Fig 4A). Using flow cytometric analysis of brain infiltrating immune cells, we observed a reduction in the priming of CD8 T cells that are specific to the immunodominant peptide VP2121-130 at 7 days post-infection (dpi) in CD11c D^b^ cKO animals (Fig 4C). Although both CD11c D^b^ cKO mice and Cre-littermates have equivalent patterns of CD45+ cell infiltration throughout the course of infection, CD11c D^b^ cKO mice exhibit disrupted CD8 T cell entry into the brain acutely. Even more strikingly, of the CD8 T cells that do enter, a lower percentage are specific for the D^b^:VP2_121-130_ epitope (Fig 4D). However, by 14 and 28 dpi the expansion of virus-specific CD8 T cells in CD11c D^b^ cKO and Cre- brains are equivalent, with nearly 80% of CD8 T cells within the brain being specific to the immunodominant D^b^:VP2121-n0 epitope in both genotypes (Fig 4C, Fig 4D). These data suggest that, while CD11c+ cells are required for a rapid CD8 T cell response to intracranial TMEV infection acutely, additional APCs are capable of priming antiviral T cells at later times. However, this response is significantly delayed relative to the rapid response mediated by CD11c+ APCs. Where CD11c D^b^ cKO animals mounted a temporally delayed antiviral CD8 T cell response against TMEV, LysM D^b^ cKO animals displayed an antiviral response that was highly kinetically similar to Cre- littermates but of a somewhat reduced magnitude. Interestingly, not only was the total CD8 T cell infiltration somewhat diminished across early stages of infection relative to what was observed in the brain tissue of Cre-littermates, but the entire CD45+ compartment exhibited a modest, but statistically significant, quantitative defect (Fig 4E). Together, these data demonstrate the required role for MHC class I on DCs for early T cell priming events while establishing a second, delayed, priming event that is DC MHC class I independent.

### CD11c D^b^ cKO mice are capable of clearing virus at later time points following infection

Because the restoration of D^b^:VP2_121-130_ specific CD8 T cells within the brain by 14 dpi in CD11c D^b^ cKO animals was striking, we next assessed the functionality of this population in clearing virus from the CNS. Because intracranial TMEV infection spreads through neuronal connections, we evaluated viral load in brains isolated at 7 dpi and spinal cords at 14 dpi (Fig 5A) [36, 37]. The number of viral plaques per gram of CNS tissue was found to be significantly higher in CD11c D^b^ cKO animals than Cre- littermates at the 7 dpi timepoint. However, by 14 dpi the CD11c D^b^ cKO animals had regained control over the virus (Fig 5B, Fig 5C). This suggests not just a quantitative but also a functional restoration of the H-2D^b^:VP2_121-130_ epitope specific CD8 T cell pool following TMEV infection.

**Fig 5:**
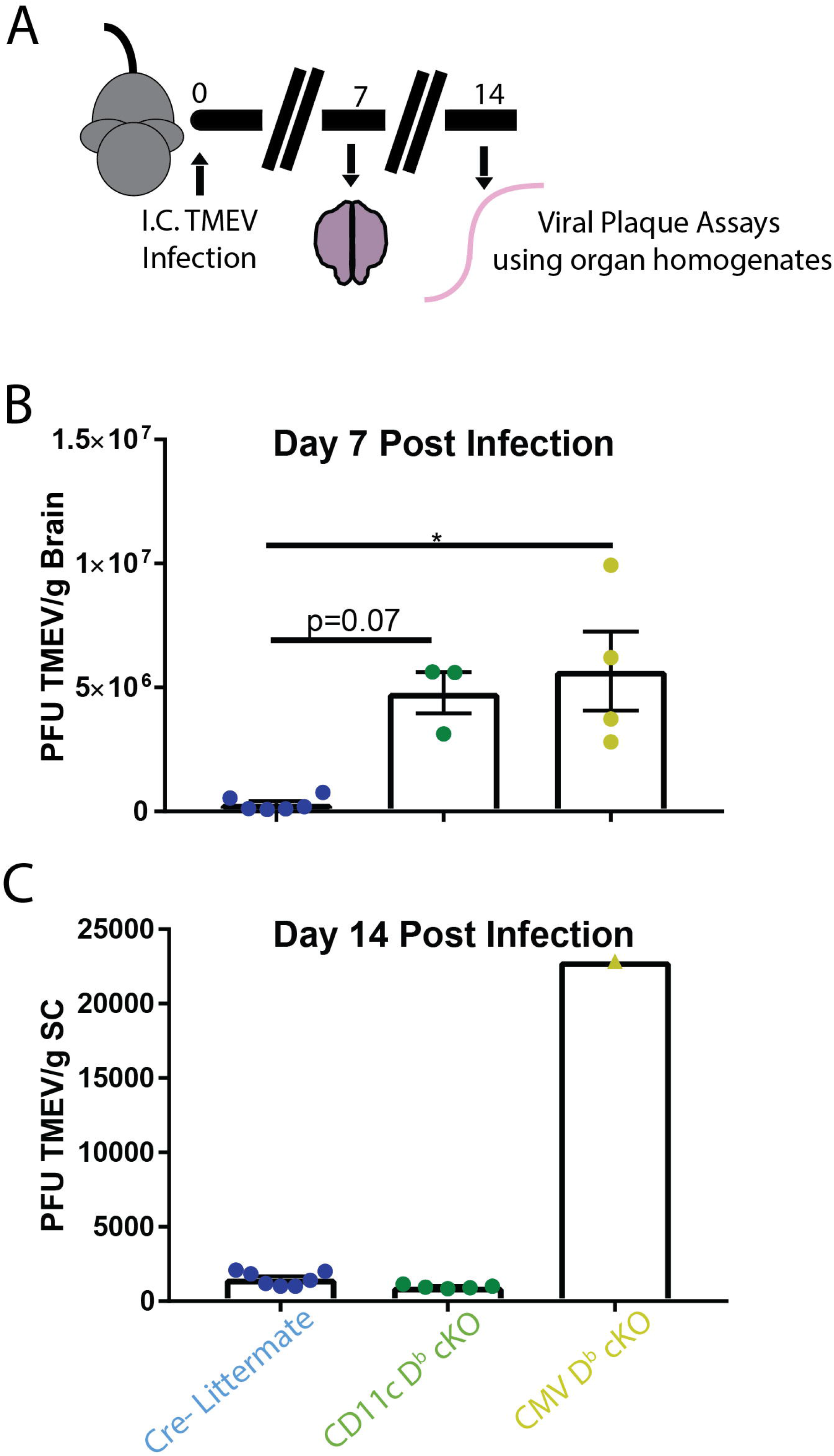
Loss of viral control observed in CD11c D^b^ cKO animals is regained by 14 dpi. (A) Experimental schematic for the analysis of CNS viral load through L2 cell plaque assay conducted with brain tissue homogenate at (B) 7 dpi and with spinal cord homogenate at (C) 14 dpi. Mice were age and gender matched within each experiment and N≥3 for each group besides 14dpi CMV D^b^ cKO. Analyses conducted with Kruskal-Wallis test. Data are presented as mean ± SEM. *, p<0.05.

### Late-arising anti-viral CD8 T cells in CD11c D^b^ cKO mice are capable of inducing blood-brain barrier disruption

In addition to mediating viral clearance, we next assessed the capacity of early versus late-primed D^b^:VP2_121-130_ epitope specific CD8 T cells to promote neuropathology in a model of blood-brain barrier (BBB) disruption developed by our research program [38-43]. This peptide induced fatal syndrome (PIFS) model is predicated on the perforin-dependent activity of previously activated CD8 T cells restimulated through intravenous administration of VP2_121-130_ peptide early in the timecourse of TMEV infection. Administration of VP2_121-130_ peptide results in extensive CNS vascular permeability, characterized by significant cerebral endothelial cell tight junction remodeling, microhemorrhage formation, neurological deficits, and death within 48 hours [38-43]. We induced PIFS through VP2_121-130_ administration in the CMV D^b^ cKO and CD11c D^b^ cKO animals, as well as Cre- Littermate controls (Fig 6A). Interestingly, at 7 dpi the CD11c D^b^ cKO animals failed to demonstrate overt symptoms at the time by which Cre- Littermates were becoming moribund (data not shown). At onset of visible symptoms in Cre- littermates, animals randomly selected from each group were subjected to T 1-weighted, gadolinium-enhanced, MRI (Fig 6A). MRI imaging revealed considerable gadolinium enhancement – consistent with CNS vascular permeability – in the brains of Cre- animals. In contrast, CD11c D^b^ cKO littermates had CNS vascular permeability indistinguishable from ubiquitous MHC-I cKO animals, who are protected from the CD8 T cell mediated BBB disruption (Fig 6B). Following MRI imaging, all animals were injected intravenously with a FITC-albumin conjugate and then euthanized one hour later, permitting analysis of CNS vascular permeability with a second methodology (Fig 6A) [44]. Measurement of fluorescence from brain homogenate after euthanasia confirmed the findings from the MRI, with increased accumulation of FITC-albumin in the brains of Cre- littermates relative to CD11c D^b^ cKO mice (Fig 6B). This leakage of FITC-albumin (green) was also visualized through immunofluorescent imaging along with the tight junction protein Occludin (red) in coronal brain sections. Cre- animals, but not CD11c D^b^ cKO littermates displayed significant disruption of tight junction integrity along the blood vessels and a diffuse spread of FITC+ regions throughout the brain parenchyma (Fig 6C). Assessment of CNS vascular permeability in the PIFS model was repeated in mice 14 dpi. At this time point, CD8 T cell responses against the immunodominant viral epitope are restored (Fig 4C). With numerically equivalent CD8 T cell responses, the capacity to induce BBB disruption was restored in CD11c D^b^ cKO mice, with regions of gadolinium enhancement and FITC-albumin accumulation both becoming equivalent to patterns observed in Cre- littermates (Fig 6D).The late-arising D^b^:VP2_121-130_ epitope specific CD8 T cell response therefore restored its potency to induce BBB disruption in the PIFS model. These data not only establish the functionality of T cells primed later in the course of infection, but also, importantly, suggest a minimum quantitative threshold of CD8 T cells required to cause pathology.

**Fig 6:**
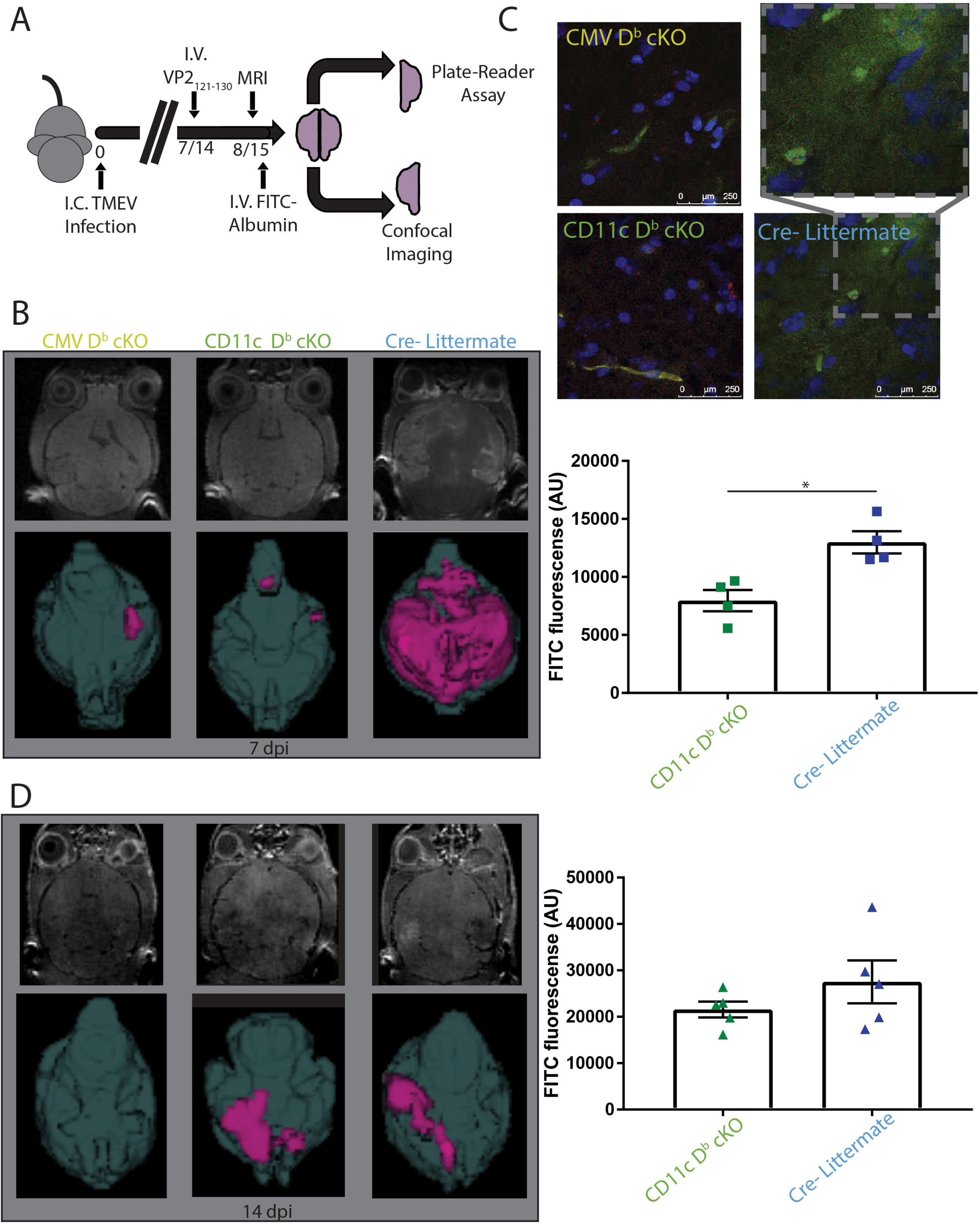
Late arising virus-specific CD8 T cells in CD11c Db cKO animals retain the capacity to elicit BBB disruption. (A) Experimental schematic for the induction of blood brain barrier disruption in mice 7 or 14 days post TMEV infection through the administration of VP2_121-130_ peptide via tail vein injection. FITC-albumin injected prior to euthanasia the following day was used along with (C) immunostaining for Occludin to visualize vessel integrity 7 dpi, and also to measure vascular leakage by assessing fluorescence intensity from brain homogenates of standardized concentration with a plate reader. Representative T 1-weighted gadolinium enhanced MRI imaging of these animals prior to euthanasia at (B) 8 dpi and (D) 15 dpi (p=0.13) are presented next to the FITC-albumin brain homogenate fluorescence data from the appropriate time point. Each experiment is conducted N≥3 for each genotype and analyses are conducted with one-way ANOVA. Data are presented as mean ± SEM. *, p<0.05.

## DISCUSSION

Studying the nature of antigen presentation in the context of CNS pathogens has traditionally required ablation of whole APC subpopulations or bone marrow transfers [45-48]. The novel H-2D^b^ cKO transgenic mouse system has laid the groundwork to study the crucial role for DCs and other APC subsets in CD8 T cell priming against CNS pathogens without disrupting other functions of these cell types, such as CD4 T cell priming or cytokine secretion. The presently described H-2D^b^ cKO mouse, in conjunction with our previously published H-2K^b^ cKO mouse, enables assessment of antigen presentation in disease models [49]. Here, we have articulated the requirement for CD11c+ cells in priming acute CD8 T cell responses against intracranial infection by the picornavirus TMEV and shown that LysM+ APCs play a subordinate role in this process. Furthermore, our data supports a role for a compensatory APC population capable of priming antiviral T cell responses, albeit much more slowly, in the absence of the antigen presentation ability of CD11c+ cells.

The importance of CD11c+ cells in the generation of a robust antiviral CD8 T cell response has been previously put forward and is further confirmed *in vivo* with the results in this study using our CD11c D^b^ cKO model [48, 50]. Using a series of bone-marrow transfer experiments, others have shown that the APC responsible for priming CD8 T cells against the VP2_121-130_ epitope in an H-2D^b^ restricted manner must be a cell of hematopoietic identity [46]. Their study supported a working model in which the initial priming of naïve CD8 T cells occurred in secondary lymphoid organs. However, it is conceivable that naïve T cells could gain entry to the inflamed brain parenchyma and be primed *in situ*; CD8 T cells displaying a naïve phenotype, and associated lack of functionality, have been previously reported to circulate through the unchallenged brain [51]. Furthermore, using an *ex vivo* organotypic culture system model, exogenous naïve CD8 T cells added to an ovalbumin (OVA) antigen-injected brain slice could enter the tissue, cluster around CD11c+ cells, and gain an activated phenotype. The ablation of CD11c+ cells in the aforementioned model was sufficient to entirely disrupt T cell priming 72 hours post injection of OVA [45]. These studies, coupled with the literature that supports the priming of T cells within tertiary lymphoid organs, are permissive of the possibility of non-canonical locations of T cell priming by CD11c+ APCs [52-54]. Considering established literature related to T cell priming, we envision four, non-mutually exclusive, contexts in which the clearly pivotal CD11c+ cells may be cross-presenting antigen: (1) DCs in secondary lymphoid organs acquire viral antigens draining out of the CNS via lymphatics, (2) rare DCs from the CNS itself are acquiring viral antigens and trafficking to draining lymph nodes, (3) unidentified migratory APCs acquire antigen and traffic to draining lymph nodes where they transfer viral antigen to CD11c+ cells, and, finally, (4) rare CD11c+ cells within the CNS are capable of stimulating CD8 T cells within the CNS, sidestepping the need for secondary lymphoid tissues for the priming process [45, 55-57]. While our novel mouse model has further established the significance of CD11c+ cells in this model of intracranial viral challenge, the migratory dynamics of involved APCs remain a topic of further investigation.

A striking result from the present study is the restoration of D^b^:VP2_121-130_ epitope-specific, functionally-antiviral, CD8 T cells in the brains of CD11c D^b^ cKO animals at 14 and 28 dpi. This rise in CD8 T cell numbers in CD11c D^b^ cKO mice, at a time point when the T cell response in control Cre- littermates is contracting, suggests a compensatory APC population that is capable of initiating a CD8 T cell response against pathogens within the CNS when CD11c+ cells are unable to effectively present antigen. The APC responsible for this delayed CD8 T cell priming remains sufficient to induce an adaptive response capable of clearing TMEV infection. Furthermore, these late-arising T cells have the capacity to induce BBB disruption in an antigen specific manner, as demonstrated in the PIFS model [58]. These observations alone do not necessarily imply that CD8 T cell populations raised by different APCs utilize the same D^b^:VP2_121-130_ epitope specific T cell receptor (TCR). It remains possible that different clonal pools of antiviral T cells exist between CD11c D^b^ cKO and Cre- animals at the 14 day time point. Analysis of these two seemingly equally functionally antiviral T cell populations for differential Vβ TCR usage is an area of future research. Finally, the identity of the APC mediating compensatory CD8 T cell priming activity in the CD11c D^b^ cKO animals remains under investigation.

One possible APC that might be responsible for the compensatory priming of D^b^:VP2_121-130_ specific CD8 T cells are MΦs. At least in other organ systems, macrophages have been suggested to play a role in naïve T cell priming [29, 59]. Bone marrow-derived, peptide-pulsed MΦs have been shown to prime CD8 T cell responses *in vivo* against a D^b^ restricted LCMV antigen, although it is interesting to note that, when peptide pulsed APCs were injected subcutaneously, MΦs needed to be exposed to three times the concentration of peptide than what was necessary for DCs in order to prime a numerically equivalent T cell response [60]. This finding could correlate with the slower increase in CD8 T cell responses in our CD11c D^b^ cKO animals, with MΦs potentially having reduced involvement in the priming of CD8 T cell responses against TMEV until a higher viral load generates sufficient levels of antigen. A study conducted by our research program in parallel to the one currently presented, utilizing the LysM-cre system in K^b^ LoxP transgenic animals, had similar results in regards to the relative importance of MΦs in CD8 T cell priming. Conditional depletion of H-2K^b^ based antigen presentation ability in MΦs resulted in no defect in the priming of CD8 T cells at 7 dpi against the K^b^:OVA epitope, which served as the immunodominant epitope in the engineered OVA-TMEV [49]. Again, similar to what has been put forward in regards to the present study utilizing the D^b^ allele and a natural viral epitope, it is possible that the contribution of any potential MΦ populations to priming this antiviral CD8 T cell response are minimal in comparison with those of CD11c+ cells and are therefore masked as long as DCs remain antigen presentation competent [49].

Microglia are another potential compensatory APC that might be responsible for the activation of D^b^:VP2_121-130_ specific CD8 T cells in our conditional knockout system. It has been previously established that activated microglia in C57BL/6 mice can cross-present soluble OVA antigen to naïve CD8 T cells *in vivo*, and other works have confirmed that TMEV infection is sufficient to drive microglia towards an activated, antigen presenting cell-like, phenotype [27, 61]. Together, these data are permissive of a role for microglia in the priming of naïve CD8 T cells against the TMEV immunodominant VP2_121-130_ epitope. On the other hand, there are conflicting data which cast doubt on the likelihood of this scenario, such as the observed abrogation of the D^b^:VP2_121-130_ epitope specific CD8 T cell response in TMEV-infected splenectomized LTα−/− animals [46]. This experiment looked only at early time points, however, so it remains unknown whether CD8 T cell priming could occur physically within the CNS at later time points during TMEV infection [46]. The ability of microglia to mediate such naïve CD8 T cell priming is currently under investigation by our research program.

In summary, adaptive immune responses, and particularly those of CTLs, are pivotal to host defense against viruses. In addition, these lymphocytes, themselves, can be drivers of neuropathology. Therefore a more robust understanding of how APCS drive this form of immunity could have profound impact on the development of effective therapies as well as limiting toxicities that arise from mobilizing activated effector CD8 T cells into the brain [62, 63]. In furtherance of these goals, we have described herein a cre-recombinase system for the cell-specific deletion of H-2D^b^ on CD11c+ cell populations in C57BL/6 mice. In the process, we have shown these APCs to be the significant driver of the acute CD8 T cell responses against TMEV infection. This work also provokes new questions about additional APC(s) responsible for CTL priming at later time points. Importantly, we introduce a novel transgenic mouse model that can serve as a critical reagent for more detailed investigations of CTL priming both within and outside the CNS moving forward.

## METHODS

### Generation and crossing of transgenic mice

Animal experiments were conducted in the manner approved by the Mayo Clinic Institutional Animal Care and Use Committee. Our group created the H-2D^b^ LoxP mouse by introducing flanking LoxP sites into the transmembrane domain of a previously cloned D^b^ transgene through site directed mutagenesis [33]. Excision of the transmembrane domain of the MHC class I molecule at the DNA level during Cre recombinase expression prevents the protein from becoming expressed on the cell surface. The Mayo Clinic Transgenic Mouse Core (Rochester, MN) was responsible for incorporating the transgene into C57BL/6 mice, who, through subsequent backcrossing with mice on an MHC-I KO background, were bred until the transgenic D^b^ was their only expressed MHC class I molecule. CD11c-Cre (B6.Cg-Tg(Itgax-cre)1-1Reiz/J, 008068) and CMV-cre (B6.C-Tg(CMV-cre)1Cgn/J, 006054) animals were purchased and crossed to MHC-I deficient C57BL/6 mice for no less than three generations to remove endogenous MHC-I molecules (The Jackson Laboratory, Bar Harbor, ME). Crossing these Cre-recombinase expressing mice with the D^b^ LoxP transgenic animals described above resulted in cKO of H-2D^b^, the presence of which could be determined by PCR for Cre using primer sequences from The Jackson Laboratory.

### TMEV infection and induction of BBB disruption

Infection and BBB disruption were conducted as previously reported [58]. Briefly, animals between 6 and 14 weeks old, were infected intracranially with 2×10^4^ PFU of Daniel’s Strain TMEV in the right hemisphere of the brain in a total volume of 10μL following anesthetization with 2% isoflurane. Mice were euthanized for flow cytometric analysis, or induced to undergo PIFS, at 7, 14, or 28 dpi. The PIFS model employed by our group involves the tail vein injection of 100μL of a 1mg/mL solution of VP2121-130 peptide (FHAGSLLVFM) in PBS at either 7 or 14 days after the original TMEV infection (GenScript, Nanjing) [38-43]. Animals were observed until they became visibly distressed (hunched and unresponsive), and then were injected via tail vein with 100μL of a 100mg/mL FITC-albumin solution in PBS (Sigma-Aldrich, St. Louis, MO). PIFS induced mice were euthanized 1hr after FITC-albumin injection for analysis.

### MRI Analysis

MRI capture and analysis was conducted as previously reported [38]. Briefly, a random subset of animals from each genotype within PIFS experiment cohorts were subjected to T1-weighted gadolinium-enhanced MRI imaging with a Bruker Avance II 7 Tesla vertical bore small animal MRI system (Bruker Biospin, Billerica) roughly 5hr before PIFS endpoint associated behaviors emerged. Mice were injected intraperitoneally with a 100mg/kg dose of gadolinium, anesthetized with 3% isoflurane and, following a 15 minute wait for gadolinium circulation, subjected to a T1-weighted spin echo sequence under maintenance dosing of 1.5% isoflurane. Their respiratory rate was monitored throughout with an MRI compatible vitals monitoring system (Model 1030, SA Instruments, Inc., Stony Brook, NY). Resultant images were processed with the Analyze12.0 software to generate object maps of regions of gadolinium enhancement. 3D models were generated by overlaying these object maps onto 20% opaque object maps of the entire brain (Biomedical Imaging Resource, Mayo Clinic, Rochester, MN).

### FITC-albumin Permeability Assay

A FITC-albumin based permeability assay was conducted as previously reported by our group [39]. In short, the left brain hemisphere taken from FITC-albumin injected animals was homogenized in RIPA buffer (50mM Tris-HCl, 150mM NaCl, 1% NP-40, 0.5% Sodium deoxycholate, 0.1% SDS, and 5mM EDTA) (Boston BioProducts, Ashalnd, MA) along with a protease inhibitor cocktail using a PowerGen 125 homogenizer (Fisher Scientific, Hampton, NH). Samples were subsequently centrifuged for 10min at 4C at 10,000rpm. The supernatant was subjected to a BCA protein assay (Pierce Biotechnology, Waltham, MA) and measured on a Synergy H1 Hybrid Multi-Mode Reader (BioTek, Winooski, VT). After normalizing the samples to the lowest sample concentration through the addition of PBS, the brain homogenates were measured on a fluorescent plate reader at 488nm excitation and 525nm emission.

### Confocal Microscopy

Confocal analyses were conducted as previously reported [49]. The right brain hemispheres taken from FITC-albumin injected animals were fresh frozen at −80°C. For analysis, they were embedded in Tissue-Tek OCT (Sakura Finetek, Torrance, CA) and were sectioned with a cryostat. 10μm coronal sections cut from near the hippocampal formation as previously described were placed on positively charged slides which were washed with PBS before fixation in 4% paraformaldehyde for 15min at room temperature. All slides were washed thrice with 100μL PBS and then incubated for an additional 1hr in 100μL of a 5% normal goat serum and 0.5% Igepal CA-630 solution (Sigma-Aldrich, St. Louis, MO). Samples were subjected to a primary stain with 100μl of a 1:200 dilution rabbit anti-mouse Occludin (clone OC-3F10, Invitrogen, Carlsbad, CA) overnight at 4C. After three washes with PBS, the Alexa Fluor 647 Goat anti-rabbit IgG secondary was applied at a 1:500 dilution in a total volume of 100μl for 1hr (Invitrogen, Carlsbad, CA). After five final washes with PBS, slides were dried and covered with Vectashield medium + DAPI (Vector lab, Burlingame, CA). Images were taken at room temperature using a Leica DM2500 with a 63x oil immersion objective, and were subsequently analyzed with Leica Acquisition Suite software (Wetzlar, Germany).

### Flow Cytometry

Flow cytometric measurements were conducted as previously published [49, 64]. Spleens and thymi harvested from euthanized animals were crushed through a 70um filter into 10mL RPMI to create a single cell suspension. Brains were harvested directly into 5mL RPMI before application of a dounce homogenizer. This brain homogenate was passed through a 70μm filter into a Percoll solution (9mL Percoll, 1mL 10x PBS, and 10mL RPMI) and subsequently centrifuged at 7840xg. After the myelin layer was removed, the remaining 10mL of brain single cell suspension was spun at 400xg for 10min along with the single cell suspensions of spleens or thymi. All samples were plated in 96-well V bottom plates and PBS washed following an ACK lysis step. If applicable, brain samples were stained with 50μL of a 1:50 dilution of D^b^:VP2121-130 tetramer (NIH Tetramer Core Facility) during a 30min incubation step in the dark at room temperature. The relevant combination of antibodies for the experimental question were subsequently applied at 1:100 dilutions during a 30min incubation on ice in the dark in 50μL total volume; these antibodies were against CD45 (V450, Clone 30-F11, Tonbo), TCRβ (PE-Cy7, Clone H57-597, Tonbo), CD8α (BV785, Clone 53-6.7, BioLegend), and CD4 (PE, Clone GK1.5, BD Pharmingen). In order to assess the usage of Vβ regions within TCRs, individual spleens were plated into 15 separate wells on a V bottom 96 well plate and each exposed for 30min on ice to 20μL of separate, pre-diluted, FITC-conjugated monoclonal antibodies in addition to a 30μL solution of 1:100 diluted antibodies against CD45 (V450, Clone 30-F11, Tonbo), TCRβ (PE-Cy7, Clone H57-597, Tonbo), CD8 (BV785, Clone 53-6.7, BioLegend), and CD4 (PE, Clone GK1.5, BD Pharmingen), as well as 1:1000 Fixable Viability Dye (eFluor780, eBioScience). Other antibodies utilized at 1:100 dilution include anti-CD11c (PerCP-Cy5.5, Clone N418, Tonbo), H-2D^b^ (FITC, Clone B22-249.R1, Accurate Chemical), F4/80 (PE, Clone BM8.1, Tonbo), B220 (APC, Clone RA3-6B2, BioLegend), Gr-1 (APC, RB6-8C5, BioLegend), and CD11b (101239 BV650, Clone M1/70, BioLegend). Viability dye (Ghost Red 780, Tonbo) at 1:1,000 was also employed for each flow cytometric experiment. Each sample was digitally compensated with single stain controls and analyzed using FlowJo v10 (FlowJo LLC, Ashland, OR).

### Plaque Assay

Viral plaque assays were conducted on L2 cells (grown in DMEM + 10% fetal bovine serum + 1% penicillin-streptomycin) with spinal cord or brain homogenates, as previously published [65]. Whole brains were homogenized in 1mL RPMI with a Dounce homogenizer and spinal cords in 1mL RPMI/100mg tissue with a PowerGen 125 homogenizer (Fisher Scientific, Hampton, NH). All homogenates were sonicated for 10s prior to application onto 12-well plates seeded with 1×10^6^ L2 cells/well. A 3% agarose overlay was added after 1hr incubation with virus, and 4 days later the cells were fixed with EAE fixative (60% Formaldehyde, 30% EtOH, 10% Glacial Acetic Acid), stained with Crystal Violet solution (1g Crystal Violet, 100mL 20% EtOH), and plaques were counted by hand.

### Statistical Analysis

All data presented as mean ± standard error of the mean (SEM), and all analyses were conducted with GraphPad Prism 7.0 (La Jolla, CA). Tests used include a two-sided Student’s *t* test, a one-way ANOVA with Holm-Sidak correction for multiple comparisons, and a Mann-Whitney Rank Sum Test if the data did not follow a normal distribution.

